# Sensorimotor basal ganglia circuit asymmetry explains lateralized motor dysfunction in early Parkinson’s disease

**DOI:** 10.64898/2026.03.16.711841

**Authors:** Elior Drori, Nitzan Kurer, Aviv A. Mezer

## Abstract

Parkinson’s disease (PD) is characterized by an early spatial pattern of degeneration in the basal ganglia. This pattern includes both posterior predominance and hemispheric asymmetry that correspond to lateralized motor symptoms. While these spatial features are well established in the striatum using dopaminergic imaging, it remains unclear whether they reflect local or circuit-level pathology across the broader basal ganglia system. It is also unknown whether widely available structural MRI can detect these spatial characteristics with comparable sensitivity. Here, we investigate the spatial pattern of basal ganglia pathology in harmonized clinical MRI data of 136 early-stage PD patients and 60 healthy controls from the PPMI database. Using gradient analyses, we identified anterior-posterior variations and selective posterior alterations in the putamen and external globus pallidus, revealing early subregional “PD hotspots”. Hemispheric asymmetry analyses across the basal ganglia showed associations with motor symptom lateralization that were most substantial in the substantia nigra, posterior putamen, and posterior external globus pallidus, indicating preferential early involvement of the sensorimotor basal ganglia circuit in lateralized PD pathology. Joint models integrating multiple regions and MRI measurements approached the explanatory power of DaTSCAN and were consistent longitudinally. In addition, they achieved robust out-of-sample cross-validated motor asymmetry prediction across visits, providing non-redundant information beyond DaTSCAN. Together, our findings demonstrate that anatomically guided spatial analysis of routine MRI reveals subregional basal ganglia pathology that underlies motor symptom lateralization in PD. This study extends spatial characterization of the basal ganglia beyond the striatum and uncovers multiple regions and data-driven posterior “hotspots” that predict motor dysfunction asymmetry. Thus, this work advances the development of MRI-based circuit-level biomarkers of PD neurodegeneration.

## Introduction

Parkinson’s disease (PD) is characterized by anatomical patterns of basal ganglia degeneration that typically manifest clinically as asymmetric motor symptoms. Two spatial properties are particularly fundamental to PD pathophysiology: hemispheric asymmetry, contralateral to the lateralization of motor impairment, and an anterior-posterior (AP) gradient of degeneration, most prominent in the putamen, where posterior subregions show the earliest and most profound dopaminergic loss. These spatial properties are well established in the sensorimotor nigrostriatal system and form a central component of contemporary models of disease progression and symptom expression^1–4^. Moreover, they have established diagnostic value^5,6^, and asymmetry has been proposed as a potential marker of variability in motor and cognitive prognosis^7^, non-motor symptom expression, and treatment-related outcomes^3,8,9^.

Currently, the primary methods to detect these spatial patterns *in vivo* are dopaminergic PET and SPECT nuclear imaging (e.g., DaTSCAN), which capture both hemispheric asymmetry and posterior-dominant striatal denervation^3,5^. While specific and sensitive, these modalities involve radiotracer exposure, are costly and not universally available, and therefore are not typically part of routine clinical evaluation. Moreover, nuclear imaging has relatively low spatial resolution, limiting subregional analysis within basal ganglia nuclei. Hence, current in vivo asymmetry measurements focus on the striatum.

Recent MRI-based studies have demonstrated that structural and microstructural imaging measures can reflect spatial aspects of PD pathology^10,11^. This body of work has shown that quantitative MRI and semi-quantitative T1w/T2w ratio capture an AP gradient in the putamen and that this gradient is altered in PD. Additionally, it found that posterior putamen (PP) asymmetry relates to both DaTSCAN asymmetry and motor symptom lateralization, indicating that MRI spatial variation contains clinically relevant information. However, these works focused primarily on the striatum and did not test whether spatial patterns and asymmetries extend across the sensorimotor and other circuits of the basal ganglia system.

This motivates several key questions. First, it remains unknown whether widely available anatomical MRI sequences, T1-weighted (T1w), T2-weighted (T2w), and proton density-weighted (PDw), can detect PD-related AP gradients and hemispheric asymmetries across multiple basal ganglia regions. This is non-trivial because reliable comparison of structural MRI intensities within and across subjects and time points requires dedicated harmonization to reduce acquisition-related variability. Second, whether nuclei beyond the putamen, including the substantia nigra (SN) and globus pallidus (GP) exhibit MRI-measurable asymmetries that relate to clinical presentation has not been systematically assessed. Third, the extent to which multiple MRI contrasts, each with different sensitivity to biophysical tissue properties, provide complementary information about these spatial features is not known. Finally, it is unclear whether MRI-derived measures from different basal ganglia regions reflect shared versus complementary aspects of motor symptom lateralization.

In this study, we address these questions by applying multi-contrast routine MRI, together with intensity harmonization and systematic multi-region asymmetry and gradient analyses, to characterize spatial markers of PD across the basal ganglia. Across a broad set of basal ganglia regions, posterior putamen (PP), posterior external globus pallidus (PGPe), and substantia nigra (SN) emerged as specific regions in which routine MRI asymmetries were substantially associated with motor symptom lateralization. This convergence on components of the sensorimotor nigro-striato-pallidal network supports a circuit-level interpretation of lateralized pathology in early PD. We further show that multi-contrast multi-region asymmetry modeling increases sensitivity to a shared underlying clinical phenomenon, and that MRI-derived asymmetries contribute unique variance beyond DaTSCAN in explaining and predicting motor asymmetry. Together, these findings demonstrate that widely available clinical MRI can capture spatial features of PD traditionally assessed with PET/SPECT, expanding the potential role of MRI-based pathology markers and suggesting that MRI-derived asymmetries may serve as a circuit-level biomarker of lateralized pathology in PD.

## Results

### Demographics

At baseline, PD (N = 136) and HC (N = 60) groups did not differ in age (Two-sample t-test, t(194) = 0.71, p = 0.48) or sex (Pearson’s χ^2^(1) = 0.17, p = 0.68). MDS-UPDRS III scores were significantly higher in PD (t(194) = 16.70, p < 10^-39^), and MoCA scores were marginally lower in PD (t(194) = -1.95, p = 0.053), with 16.2% of PD subjects showing mild cognitive impairment. PD characteristics are shown in **Table *1***.

**Table 1.** Demographics and PD characteristics at baseline.

|  | PD (N = 136) | HC (N = 60) | Test |
| --- | --- | --- | --- |
| Age, years | 61.1 ± 9.3 (range, 38.5-82.2) | 60.0 ± 10.8 (range, 32.3-79.8) | $t(194) = 0.71$, $p = 0.48$ |
| Sex, female, n (%) | 48 (35%) | 23 (38%) | $\chi^2(1) = 0.17$, $p = 0.68$ |
| Disease Duration, years | 0.6 ± 0.6 (range, 0.0-3.0) | NA | |
| H&Y scale | 1.6 ± 0.5 (range, 1-2) | NA | |
| NSD, n (%) | 121 (89%) | NA |  |
| NSD stage | 2.7 ± 0.8 (range, 0-4) | NA | |
| MDS-UPDRS III, score | 21.0 ± 9.4 (range, 4-43) | 0.6 ± 1.4 (range, 0-8) | $t(194) = 16.70$, $p = 2e-39$ |
| MoCA, score | 27.7 ± 2.1 (range, 20-30) | 28.2 ± 1.1 (range, 27-30) | $t(194) = -1.95$, $p = 0.053$ |
| Normal (>25) | 114 (84%) | 60 (100%) |  |
| MCI (>17) | 22 (16%) | 0 (0%) |  |

H&Y: Hoehn and Yahr; NSD: Neuronal Synuclein Disease; MCI: Mild cognitive impairment

### Data Harmonization

Harmonization substantially reduced MRI intensity variability across visits and subjects. Before harmonization, whole-brain histograms for T1w, T2w, and PDw images showed poor overlap both within subjects across four visits (**Figure 1 A**) and between subjects across eight imaging centers (**Figure 1 B**). Harmonization markedly increased histogram overlap for all contrasts (Dice coefficients of HC subjects at baseline: T1w = 0.35→0.70, T2w = 0.11→0.76, PDw = 0.01→0.71), indicating improved consistency of brain intensity distributions. These improvements enabled a better comparison of regional MRI intensities across sessions and subjects in subsequent analyses.

**Figure 1.**
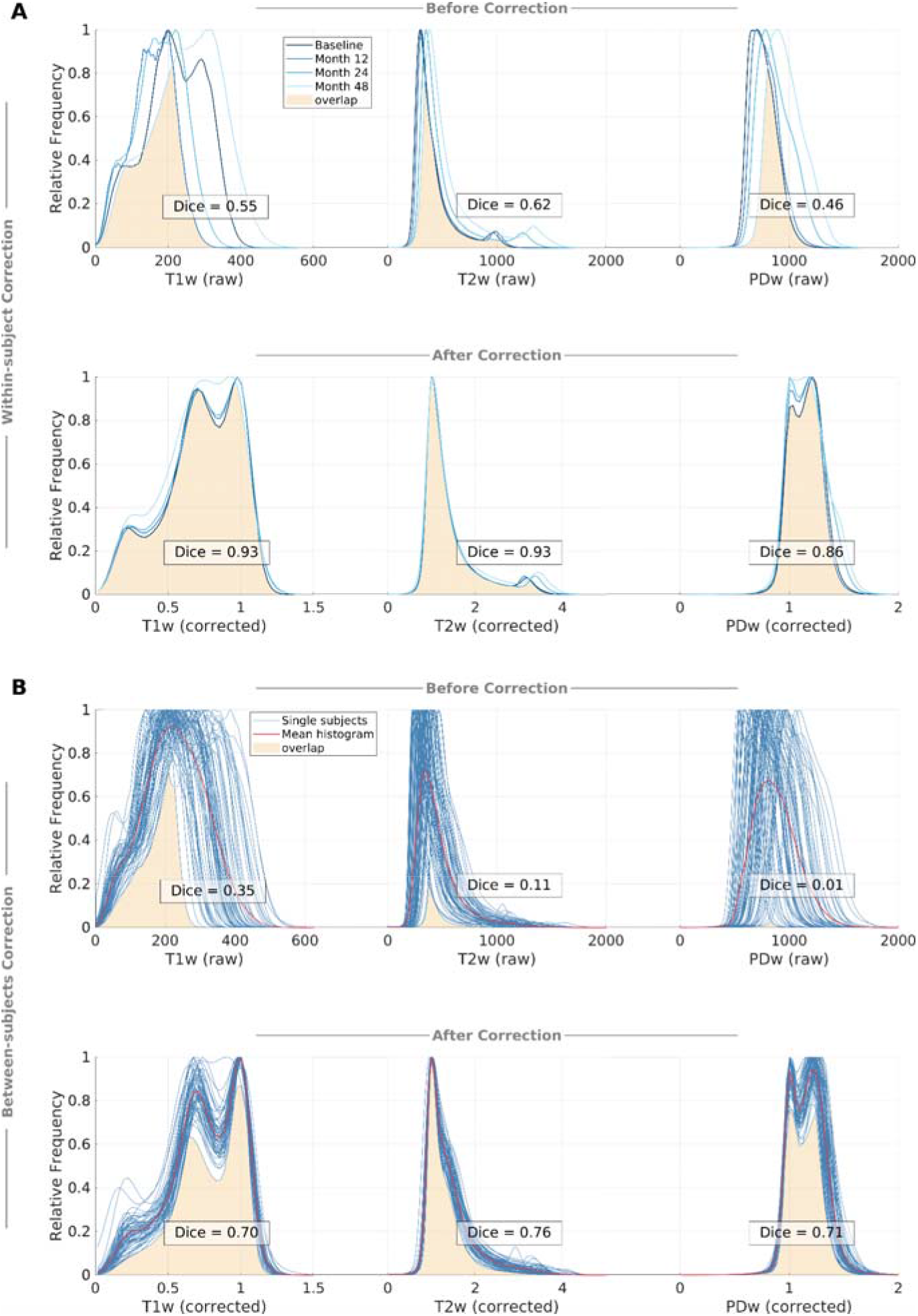
Data harmonization reduces within- and between-subject whole-brain variability. **(A)** Within-subjects harmonization: T1w, T2w, and PDw histograms of a single example HC subject show substantially improved overlap across four visits after data harmonization (Dice: T1w = 0.55→0.93, T2w = 0.62→0.93, PDw = 0.62→0.86). **(B)** Between-subjects harmonization T1w, T2w, and PDw histograms of all HC subjects at Baseline visit (N = 60), show substantial improvement after harmonization (Dice: T1w = 0.35→0.70, T2w = 0.11→0.76, PDw = 0.01→0.71).

### Putamen and GPe Multi-Contrast Gradients Alter in PD

Following earlier findings of putamen AP gradients^10,11^, we found putamen AP gradients of T1w, T2w, and PDw at the baseline visit (**Figure 2 A–C**), indicated by significant main effects of Position in the linear mixed-effects models (LMMs; see **Methods**, and **Supplementary Table 1**). We also found PD-related changes in these gradients, indicated by significant Position × Group interaction effects. Post hoc analyses (see **Methods**) localized the sources of these interactions in posterior segments of the putamen across all contrasts, highlighting the PP as a subregion preferentially affected in PD (**Figure *2* D** and **Supplementary Table 2**).

**Figure 2.**
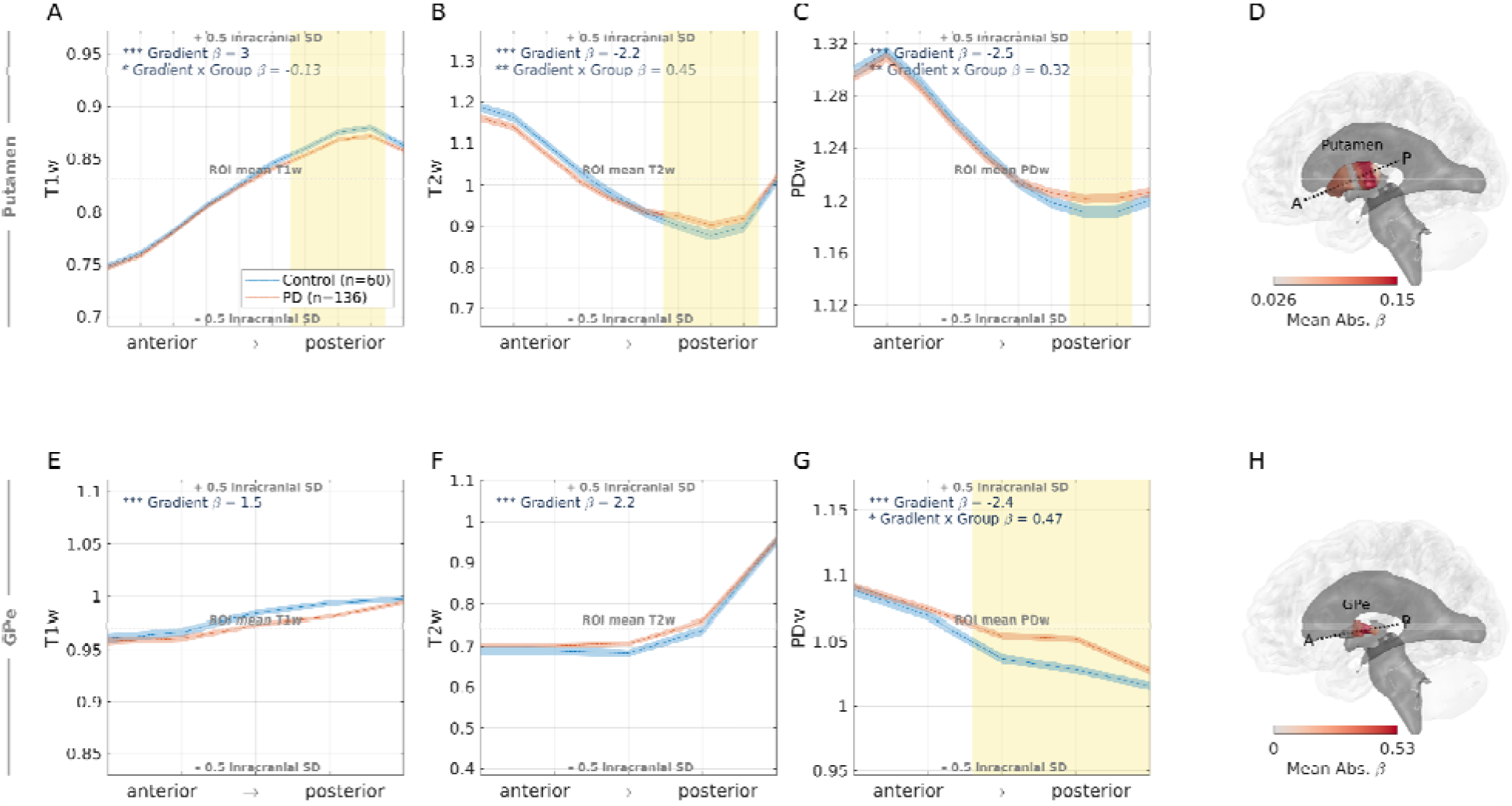
Putamen and GPe AP gradients are altered in posterior segments in PD. **(A-C)** Profiles of (A) T1w, (B) T2w, and (C) PDw intensities along the AP axis of the putamen show significant spatial variation (i.e., gradient) and significant PD-related interactions. Post hoc analyses localized the source of the PD-related change in posterior positions in all MRI contrasts (yellow shading, see **Methods**). **(D)** A summary of the localization analysis is visualized on the putamen, with mean absolute beta coefficients of AP positions across all significant contrasts highlighting the PP subregion as a PD hotspot (see **Methods**) **(E-G)** Profiles along the AP axis of the GPe show significant gradients in all MRI contrasts, and a significant PD-related change in the PDw gradient. **(H)** A summary of the localization analysis in the GPe, highlighting the PGPe as a PD hotspot. The ranges of all y-axes in this figure are defined as the ROI mean value ± 0.5 SD of the MRI contrast in the entire brain, in order to give sense of the measured variation. Data reflect the more affected side (MAS) values for PD subjects with motor asymmetry, and averaged values across left and right hemispheres for HC and PD subjects without motor asymmetry. * p < 0.05, ** p < 0.0001, *** p < 10^-27^ (FDR corrected; **Supplementary Table 1**).

Extending the gradient analysis beyond the putamen, in the GPe, we similarly identified AP gradients in all three contrasts (**Figure *2* E–G**). A PD-related change was found in the PDw gradient, with post hoc analysis localizing this effect to a broad posterior portion of the GPe (PGPe; **Figure 2 H)**.

Given prior evidence indicating no gradient changes in the caudate at early PD stages in an overlapping cohort^10^, we did not extend the gradient analysis to the caudate in order to limit additional statistical comparisons. We also did not apply this approach to smaller nuclei such as the GPi, SN, nucleus accumbens, and STN, where small volumes relative to the MRI resolution make a whole-ROI approach more reliable (see **Table 2** and **Methods**).

**Table 2.**
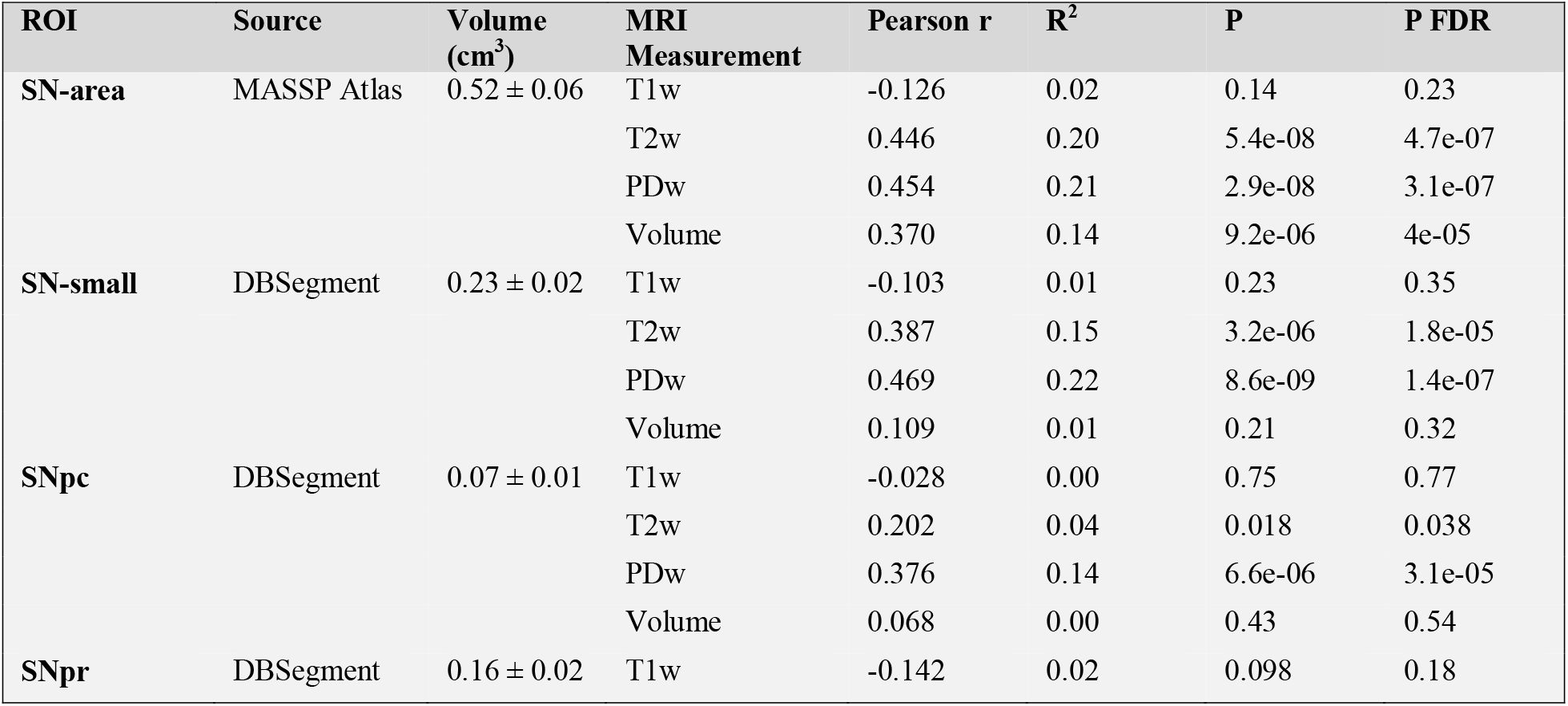
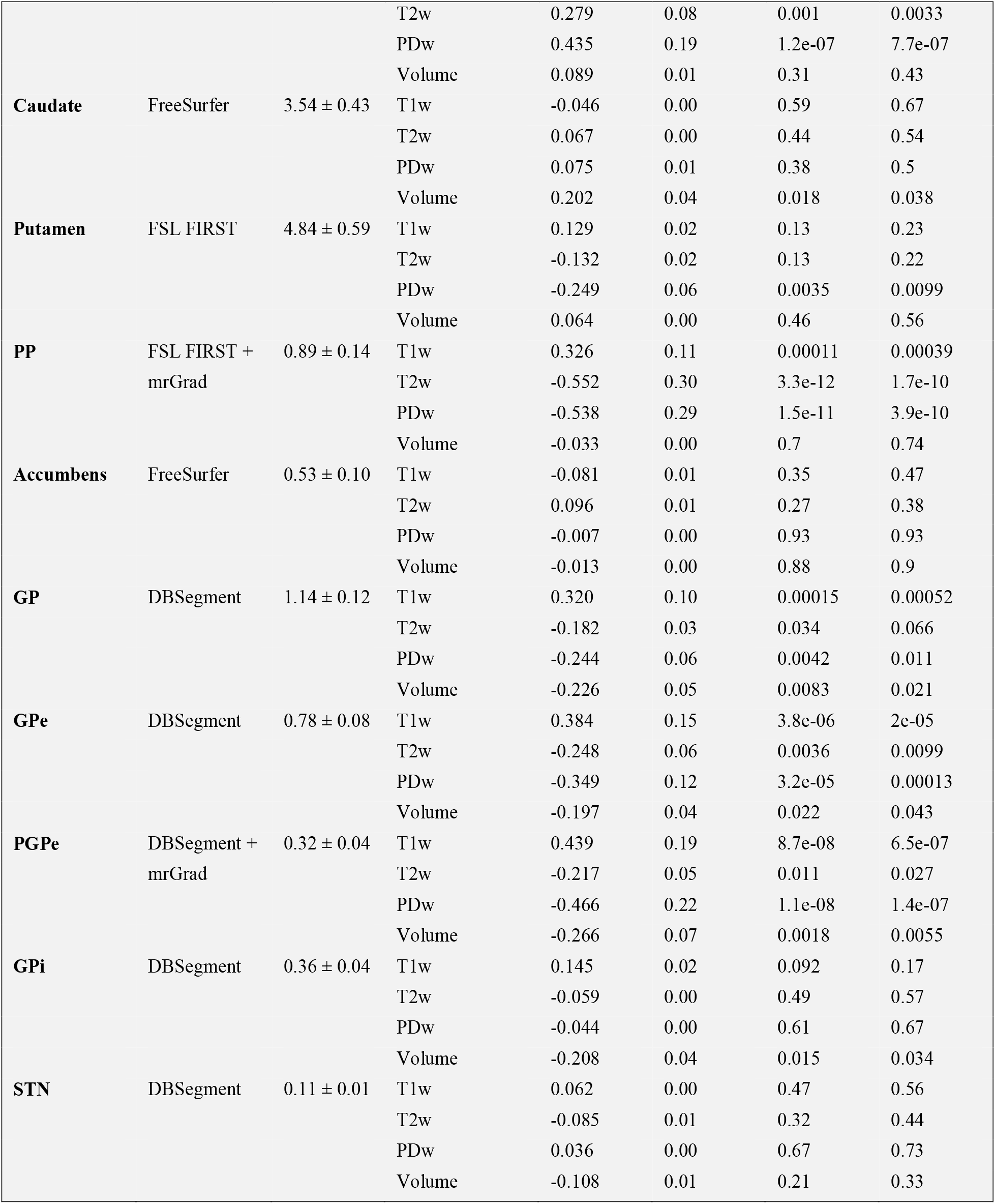
Correlations of basal ganglia MRI asymmetries with motor asymmetry in PD. Statistically significant effects after FDR correction are shown in bold.

### Basal ganglia asymmetries detected with standard anatomical MRI are associated with motor symptom asymmetry

Extending prior findings demonstrating the clinical relevance of PP structural asymmetry^10,11^, we tested the linear relationship at the baseline visit between motor asymmetry of PD patients and MRI intensity asymmetry in multiple basal ganglia regions.

ROIs included the gradient analysis-driven subregions PP and PGPe, as well as the whole SN (SN-area, identified by MASSP atlas; see **Methods**), another smaller SN ROI (SN-small, identified by DBSegment; see **Methods**) and the SN subregions: pars compacta (SNpc) and pars reticulata (SNpr), the caudate, putamen, nucleus accumbens, GP and its subregions: GPe and GPi, and STN. We found significant associations between MRI asymmetry indices and motor asymmetry across multiple basal ganglia regions, after FDR correction for multiple comparisons (**Table 2**). Effect sizes were most substantial in SN-area, PP, and PGPe. In the SN-area, significant associations with motor asymmetry were found for T2w (R^2^ = 0.20, p < 10^-7^) and PDw (R^2^ = 0.21, p < 10^-7^), but not T1w (R^2^ = 0.02, p = 0.23; all p values are FDR corrected; **Figure 3, left**). In the PP, asymmetries from all three contrasts were significantly correlated with motor asymmetry, in opposite directions compared with the SN (T1w: R^2^ = 0.11, p < 0.001; T2w: R^2^ = 0.30, p < 10^-10^; PDw: R^2^ = 0.29, p < 10^-10^; **Figure 3, middle**). In the PGPe, significant associations were observed for T1w (R^2^ = 0.19, p < 10^-7^) and PDw (R^2^ = 0.22, p < 10^-7^) and a smaller effect was observed for T2w (R^2^ = 0.05, p < 0.05; **Figure 3, right**).

**Figure 3.**
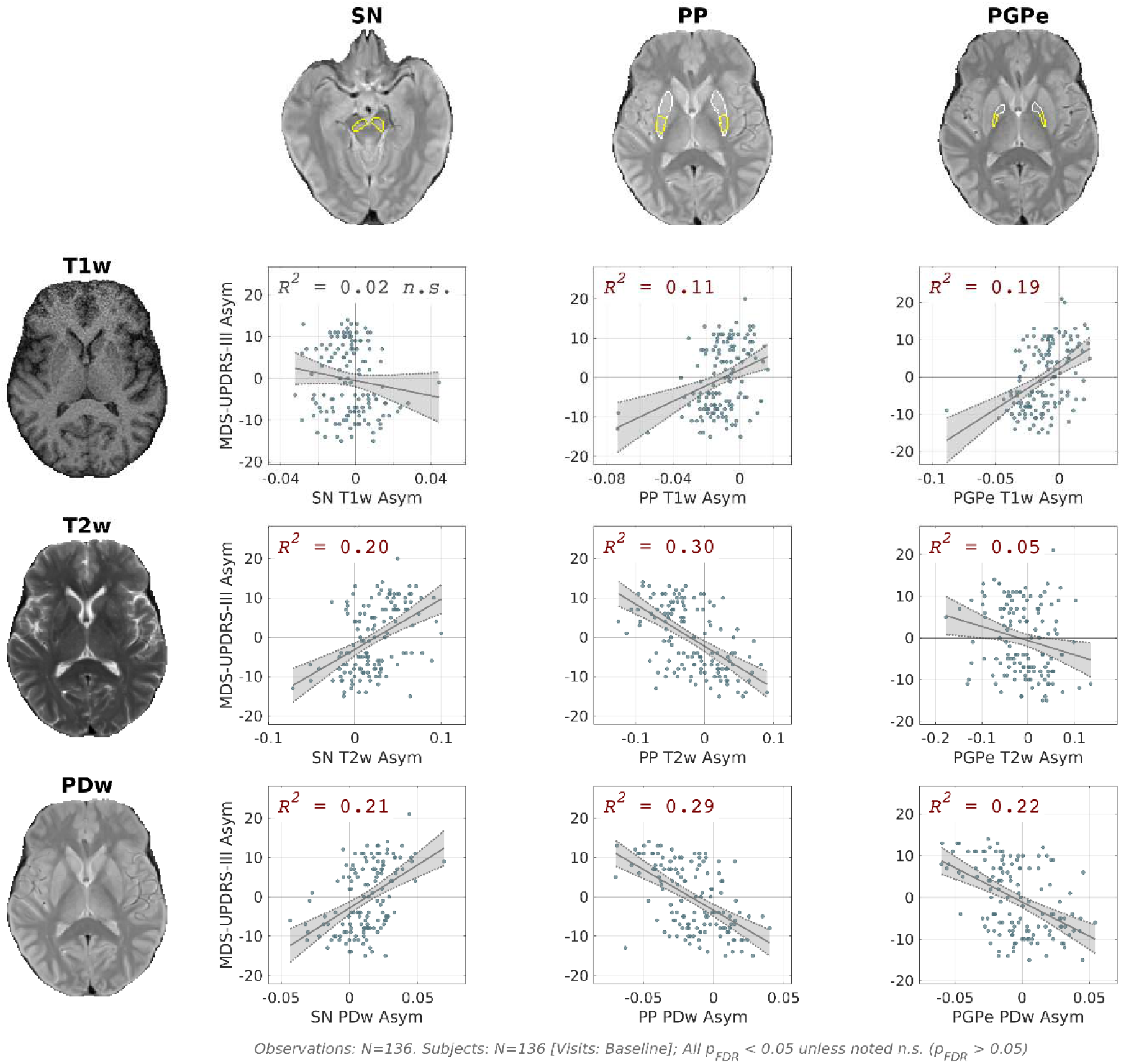
Basal ganglia MRI-derived asymmetries are associated with motor symptom asymmetry. Panels display scatter plots showing linear relationships between MRI intensity asymmetries and MDS UPDRS-III motor asymmetry in PD patients across MRI contrasts (T1w, T2w, PDw; rows) and basal ganglia ROIs (PP, SN, and PGPe; columns). Significant associations were found for PP and PGPe across all contrasts, and for SN in T2w and PDw with opposite directionality relative to PP and PGPe. Regression lines represent best-fit models; shaded areas indicate 95% confidence intervals; reported R^2^ values reflect the proportion of motor asymmetry variance explained by the MRI asymmetries. N = 136 (Baseline visit). All p_FDR_ < 0.0005, unless noted n.s. (p_FDR_ > 0.05).

In addition to these three ROIs, significant but generally weaker associations with motor asymmetry were observed in anatomically related regions. The SN-small showed significant effects for T2w (R^2^ = 0.15, p < 10^-5^) and PDw (R^2^ = 0.22, p < 10^-7^). Within SN-small, both SNpc and SNpr showed significant effects for T2w and PDw contrasts (SNpc: T2w R^2^ = 0.04, p < 0.05, PDw R^2^ = 0.14, p < 10^-5^; SNpr: T2w R^2^ = 0.08, p < 0.01, PDw R^2^ = 0.19, p < 10^-7^). Notably, the larger effect sizes in SN-area compared with the narrower SN definitions may reflect greater overlap with the substantia nigra, which is difficult to segment reliably^12^. Compared with the PGPe, weaker associations were detected in the whole GPe across contrasts (T1w R^2^ = 0.15, p < 10^-5^, T2w R^2^ = 0.06, p < 0.01, PDw R^2^ = 0.12, p < 0.001) and in the whole GP (T1w R^2^ = 0.10, p < 0.001; PDw R^2^ = 0.06, p < 0.05). Finally, compared with the PP, a weaker association with motor asymmetry was observed in the whole putamen (PDw: R^2^ = 0.06, p < 0.01).

In addition to MRI intensity asymmetries, we also tested the association of volumetric asymmetries with motor asymmetry. In the SN-area, the volume asymmetry significantly explained motor asymmetry (R^2^ = 0.14, p < 10^-5^). Additional basal ganglia ROIs showed significant but smaller volumetric asymmetry effects: caudate (R^2^ = 0.04, p < 0.05), GP (R^2^ = 0.05, p < 0.05), GPe (R^2^ = 0.04, p < 0.05), GPi (R^2^ = 0.04, p < 0.05), and PGPe (R^2^ = 0.07, p < 0.01). In regions where both intensity and volume associations were present, volumetric associations were smaller, indicating that intensity asymmetries were not explained by volumetric differences. Caudate and GPi, however, showed only volumetric associations.

Beyond the basal ganglia ROIs, an exploratory analysis examined correlations between MRI contrasts asymmetries and motor asymmetry across a broad set of cortical and subcortical regions (**Supplementary Section 1**). Several regions, including the ventral tegmental area (VTA), pedunculopontine nucleus (PPN), cerebellum, and occipital regions, showed small effect sizes; however, none of these associations remained significant after correction for multiple comparisons (**Supplementary Figure 1**).

Based on these results, subsequent analyses focused on the broad SN-area (hereafter: SN, for simplicity) and the gradient-derived subregions PP and PGPe.

### MRI measurements provide complementary information about basal ganglia asymmetry

Focusing on SN, PP, and PGPe, we tested whether combining multiple MRI contrast and volume asymmetries provides complementary information within ROI by comparing a multivariate MRI model to the best single-measurement model. For each ROI separately, we fitted a multiple regression asymmetry model that included the MRI measurements that were most informative in the single-measurement analyses (R2 > 0.10; SN: T2w, PDw, volume; PP: T1w, T2w, PDw; PGPe: T1w, PDw; see **Table 2**). We found that the multivariate models explained significantly more variance in motor asymmetry than the best single-measurement models alone (SN: ΔR^2^ = 0.13; PP: ΔR^2^ = 0.11; PGPe: ΔR^2^ = 0.10; all FDR-corrected p < 10^-5^; **Figure 4 A-C**). This suggests complementary contributions of the informative MRI measures within these regions. Moreover, within each ROI multivariate model, all included variables showed significant independent contributions (**Supplementary Table 3**).

**Figure 4.**
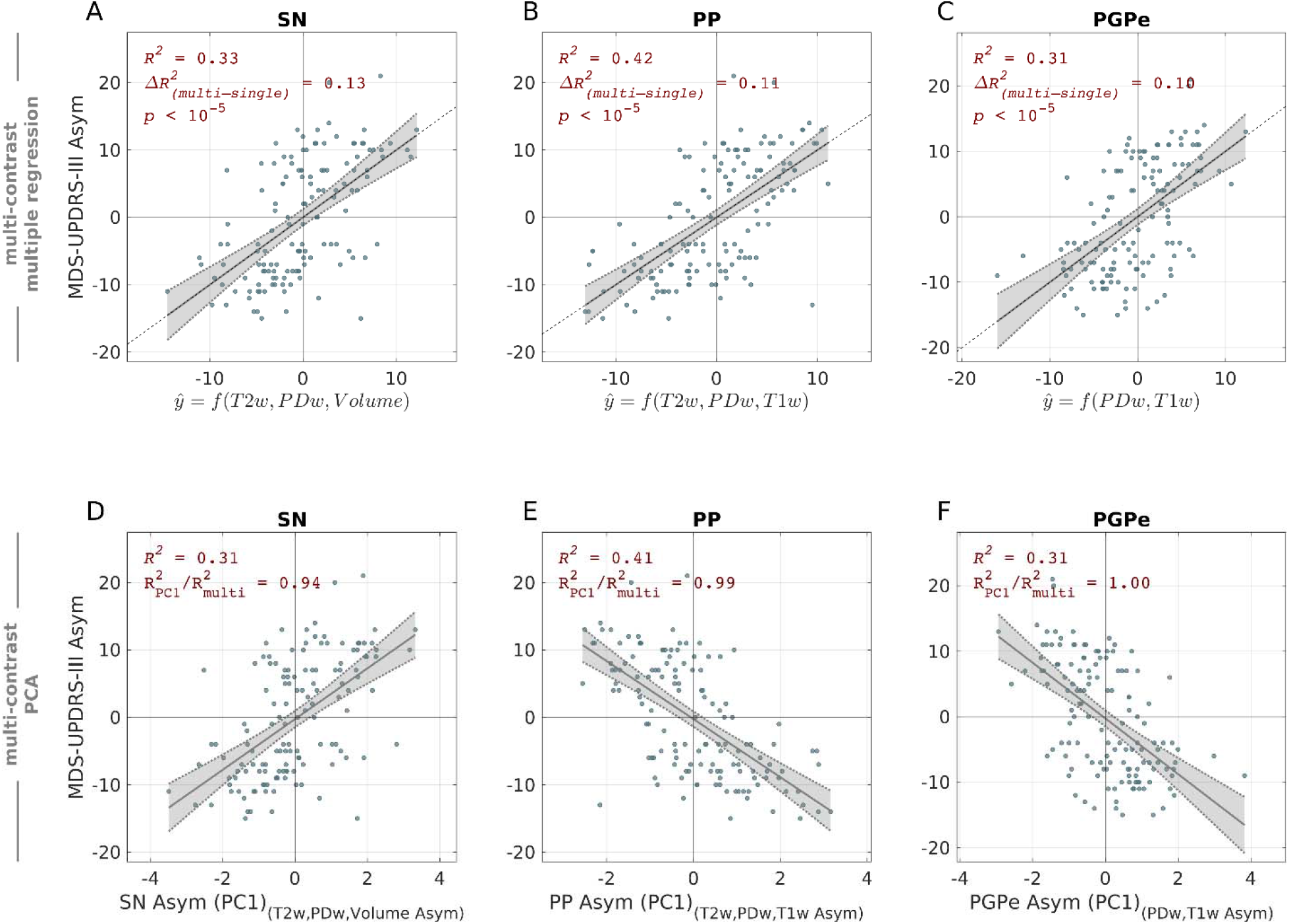
Complementary and shared MRI contributions to motor asymmetry within basal ganglia regions. **(A-C)** Multivariate regression models of motor asymmetry within ROI, using multiple MRI measurements that were found informative in previous single-measurement analyses (R^2^ > 0.10; SN: T2w, PDw, volume; PP: T1w, T2w, PDw; PGPe: T1w, PDw; see **Table 2**). Models explained significantly more variance in motor asymmetry than the best single measurement within SN (A), PP (B), and PGPe (C). In each ROI model, all included MRI measures showed significant contributions (**Supplementary Table 3**). ΔR^2^ denotes the increase in variance explained relative to the best single-measurement model (see **Figure 3**). P values indicate significance of model improvement (Nested F-test). **(D-F)** Simplified models using only PC1 (denoted SN_Asym_, PP_Asym_, and PGPe_Asym_) derived from the corresponding MRI measurements in (A-C). The variance in motor asymmetry explained by each ROI’s PC1 (capturing 56%-65% of the measurements variance) was comparable to that of the corresponding multivariate MRI model, accounting for 94-100% of its explained variance (R^2^_PC1_ / R^2^_multi_). This indicates that most of the clinically relevant variance among the selected MRI measurements within ROI is captured by a shared latent component.

To assess whether informative MRI measures reflect distinct or shared latent components within each ROI, we performed PCA on the MRI measurements included in the multivariate models. We found that the first principal component (PC1) within each ROI (termed SN_Asym_, PP_Asym_, and PGPe_Asym_) explained a comparable proportion of variance in motor asymmetry to the corresponding multivariate MRI model, with R^2^_PC1_ / R^2^_multi_ > 0.94 (**Figure 4 D-F**). This indicates that within SN, PP, and PGPe, a shared MRI component captures most clinically relevant variance despite the presence of statistically significant independent contributions.

### Multi-ROI multi-contrast MRI–motor asymmetry associations and longitudinal consistency

Next, we tested whether the selected ROI composite asymmetries SN_Asym_, PP_Asym_, and PGPe_Asym_ provide complementary information about motor asymmetry. A multiple regression model including these three composite asymmetries at baseline explained more variance in motor asymmetry than any single ROI_Asym_ (R^2^ = 0.56, ΔR^2^ = 0.15, p < 10^-9^); **Figure 5 A and Supplementary Table 4**), and all ROIs showed significant unique contributions to the explained variance.

**Figure 5.**
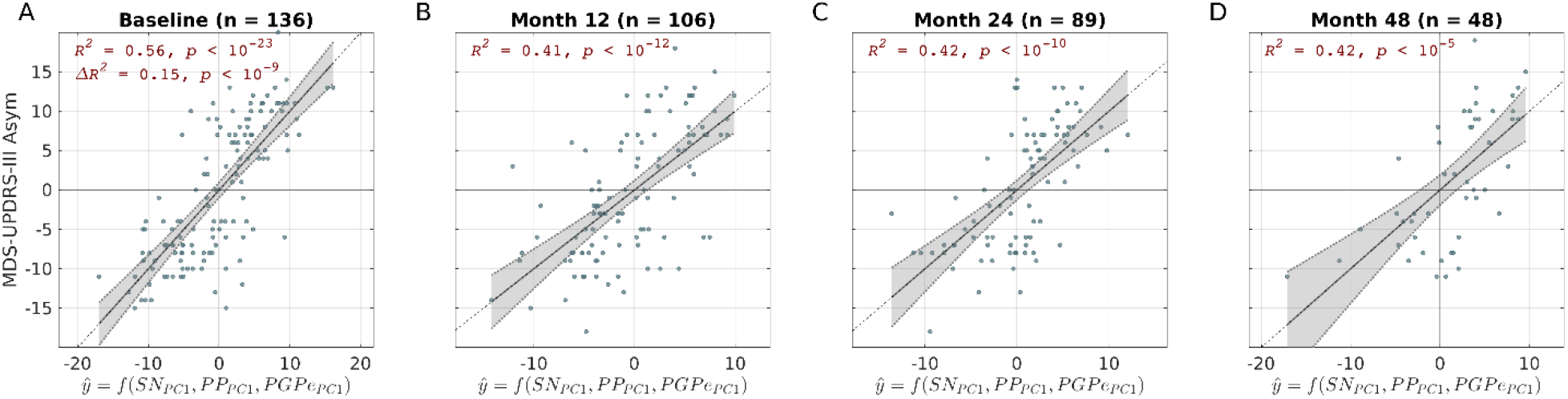
Multi-region MRI model explains motor asymmetry at baseline and follow-up. **(A)** A multi-region motor asymmetry model, combining SN_Asym_, PP_Asym_ and PGPe_Asym_, at baseline. Each ROI_Asym_ variable represents the first PC of multiple MRI measurements within that ROI (see **Figure 4 D-F**). The multi-region model explained substantially more variance in motor asymmetry than any single region alone (Δ**R**^**2**^ **= 0.15, p < 10**^**-9**^). **(B-D)** The multi-region models at 12-, 24-, and 48-month follow-up visits were evaluated using PCA loadings defined at baseline to assess longitudinal consistency. Associations observed at baseline were largely consistent in follow-up, although effect sizes (R^2^) decreased.

This multi-region model explained motor asymmetry to a degree comparable to DaTSCAN asymmetry measured in the same cohort (R^2^ = 0.58; **Supplementary Table 5**), indicating that routine MRI captures clinically relevant information typically assessed using nuclear imaging.

To assess longitudinal consistency, we evaluated the same multi-region model at follow-up visits (Month 12, 24, 48), using the PCA loadings defined for each ROI at baseline. The association observed at baseline remained largely consistent across follow-up visits, albeit with attenuated effect sizes (**Figure 5 B-D**). At Month 48, significant contributions persisted for PP and SN but were no longer significant for the PGPe.

### Predicting motor asymmetry from multi-region, multi-contrast MRI

Beyond descriptive associations within individual visits, we evaluated whether the predefined ROI asymmetries SN_Asym_, PP_Asym_, and PGPe_Asym_ predicted motor asymmetry across all visits. In a model combining the three composite asymmetry measures, repeated subject-wise Monte Carlo cross-validation (30 repetitions) yielded out-of-sample performance of *R*^*2*^_*cv, test*_ = 0.39 ± 0.14 and RMSE_*cv, test*_ = 6.25 ± 0.75 (mean ± SD across repetitions). Averaging predictions across repetitions yielded global *R*^*2*^_*cv, test*_ = 0.43 and global RMSE_*cv, test*_ = 6.21 (**Figure 6 A**). In each repetition, normalization and PCA loadings were recomputed on the training data only, whereas the ROI and MRI-measurement sets were predefined from the baseline analyses. A subject-level permutation test confirmed the statistical significance of the models (500 permutations, p < 0.002), conditional on the prior predictor selection.

**Figure 6.**
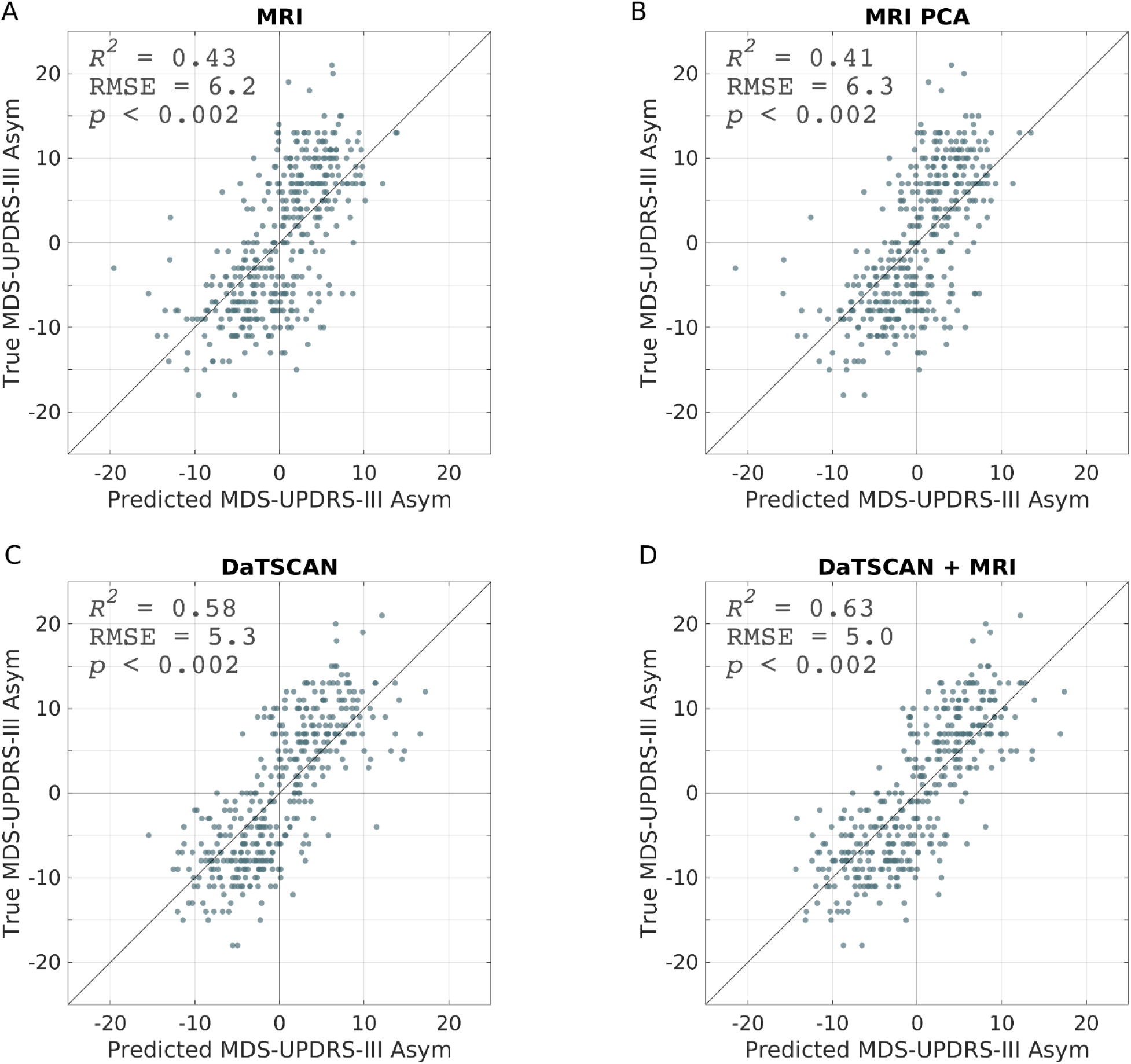
Cross-validated prediction of motor asymmetry. Scatter plots of predicted versus observed motor asymmetry, using (A) multi-region multi-contrast MRI asymmetry model; (B) multi-region multi-contrast PCA MRI asymmetry model; (C) DaTSCAN anterior putamen SBR asymmetry model; and (D) combined DaTSCAN and MRI asymmetry model. Reported values correspond to global cross-validated R^2^ and subject-level permutation p value; line = identity.

Importantly, when performing a single PCA across all selected ROIs and MRI measurements, PC1 yielded similar cross-validated performance (*R*^*2*^_*cv, test*_ = 0.38 ± 0.15 and RMSE_*cv, test*_ = 6.34 ± 0.81; global *R*^*2*^_*cv, test*_ = 0.41 and global RMSE_*cv, test*_ = 6.30; **Figure 6 B**). Across repetitions, PC1 accounted for a mean of 32.9% of the total variance in the predictors (range: 28.0 - 36.7%). This result indicates that the predictive signal was largely captured by variance shared across SN, PP, and PGPe, consistent with a circuit-level latent component.

Notably, DaTSCAN asymmetry achieved higher predictive performance than MRI (R^2^_cv, test_ = 0.58, ΔR^2^ = 0.15; **Figure 6 C**). However, adding MRI to the DaTSCAN model increased the explained variance by an additional ∼4% (ΔR^2^ = 0.04, p < 0.002), indicating that MRI contributes unique information beyond DaTSCAN nuclear imaging (**Figure 6 D** and **Table *3***).

**Table 3.** Comparison between MRI and DaTSCAN in motor asymmetry prediction.

| Modality | ROI | R<sup>2</sup> | RMSE | Modality's $\Delta R^2$ |
| --- | --- | --- | --- | --- |
| **DaTSCAN** | Ant. Putamen | 0.58 | 5.4 | 0.20 |
| **MRI** | PP, SN, PGPe | 0.43 | 6.2 | 0.04 |
| **DaTSCAN + MRI** |  | 0.63 | 5.1 | Shared Variance: 0.39 |

R<sup>2</sup> values are rounded to two decimals; $\Delta R^2$ values are computed from full-precision estimates.

## Discussion

This study shows that spatially informed analyses of widely available clinical MRI reveal hallmark spatial characteristics of PD and yield clinically meaningful information about both disease presence and the lateralized expression of motor dysfunction. We demonstrate that two spatial features of PD pathology - posterior predominance and hemispheric asymmetry associated with the clinically more affected side - can be detected using conventional T1w, T2w, and PDw images when appropriately harmonized across imaging centers and longitudinal visits. Extending prior work focused primarily on the striatum^10,11^, we further show that spatial variation within basal ganglia nuclei is non-uniform and that MRI-derived asymmetries in these structures reflect clinical motor asymmetry. These MRI asymmetry measures explained a substantial proportion of the variance in motor asymmetry, approached the explanatory power of DaTSCAN, contributed additional predictive information when added to DaTSCAN, and were longitudinally consistent. Together, these findings indicate that routine MRI, when analyzed with anatomically guided and spatially resolved approaches, can capture subregional basal ganglia pathology in vivo, and support the development of both localized and system-level biomarkers of motor dysfunction in PD.

A central finding of this study is that AP gradients within basal ganglia nuclei contain disease-relevant information. In the putamen, gradient analysis revealed PD-related changes confined to posterior subregions (PP), replicating and extending previous findings obtained with quantitative and T1w/T2w MRI^10,11^. Prior studies have suggested functional and connectivity-based AP gradients in the GP^13–16^. Here, we show that gradients in the GPe are detectable using routine MRI and that they alter in PD in posterior subregions (PGPe).

This spatially specific pattern of posterior putaminal and pallidal “PD hotspots” is in agreement with previous findings of preferential early degeneration in posterior striatal and pallidal regions in PD^1,16,17^. Moreover, identification of reproducible striato-pallidal “hotspots” may provide a principled anatomical framework for refining circuit-targeted therapeutic strategies, including neuromodulation and other regionally specific interventions^18^.

Beyond group-level spatial patterns, SN, PP, and PGPe emerged as key regions in which MRI asymmetry indices were strongly associated with individual differences in clinical motor lateralization. Importantly, these associations were anatomically specific: not all basal ganglia regions showed comparable relationships, and exploratory analyses outside the basal ganglia did not yield substantial effects. While the relevance of nigrostriatal asymmetries to PD is well established^1,4,5^, this study provides evidence that pallidal asymmetry is also directly linked to clinical asymmetry, using widely available MRI. Previous studies using arterial spin labeling MRI reported no significant group differences in pallidal asymmetry, but a direct subject-level association with motor symptom lateralization had not been examined^19,20^. By demonstrating such relationships in the GPe and its posterior subregion, our results expand the set of basal ganglia structures implicated in lateralized PD pathology and highlight the value of subregional analyses beyond the striatum.

Together, these results provide evidence of early involvement of SN, PP, and PGPe in PD, supporting contemporary theories emphasizing early, selective vulnerability of distinct neuronal populations within the sensorimotor nigro-striatopallidal circuit of the basal ganglia^2,21,22^.

Our findings further indicate that MRI contrasts provide complementary information about basal ganglia asymmetry. T1w, T2w, and PDw contrasts showed region-specific associations with motor asymmetry, consistent with their partly differential sensitivity to tissue properties such as iron content, water concentration, and atrophy-related changes^23^. Decreased T2w signal in the more affected SN (contralateral to the more affected body side) may reflect iron accumulation^24^, whereas increased T2w together with decreased T1w in the more affected PP is consistent with atrophy-related increased water content^11^. The relatively small PD-related T2w change in the GPe may reflect early-life saturation of pallidal iron concentrations^25^, limiting sensitivity to further disease-related change, as previously suggested^26^. Multiple regression showed that contrasts retained statistically detectable unique associations within ROIs, but PCA analyses indicated that most clinically relevant variance within each ROI could be captured by a shared component across informative contrasts. Thus, the data support a shared underlying clinical variance that is captured by the different MRI measurements.

While intensity asymmetries likely reflect microstructural tissue changes, volumetric asymmetry analyses indicated additional but weaker macroscopic effects. Associations between volume asymmetry and motor asymmetry were observed in several basal ganglia regions, including SN, caudate, GP, GPe, GPi and PGPe, but effect sizes were consistently smaller than those of the MRI contrast intensities. Consistent with a prior meta-analysis indicating that volumetric changes are subtle in early PD^27^, these findings support the greater sensitivity of MRI intensity measures to early tissue alterations.

Beyond complementary information within ROIs, complementary information was also observed across basal ganglia regions. A joint model including SN, PP, and PGPe composite asymmetry measures explained motor asymmetry to a degree comparable to DaTSCAN. This result supports the view that PD motor symptoms reflect dysfunction across interconnected basal ganglia nodes^28^. The MRI-motor asymmetry associations observed at baseline remained consistent at 12-, 24-, and 48-month follow-up visits, although effect sizes attenuated over time, particularly for the PGPe. This attenuation likely reflects a combination of the reduced clinical variance, decreasing sample size, and increasing heterogeneity introduced by symptomatic treatment during follow-up visits.

Importantly, MRI asymmetry indices did not only correlate with motor asymmetry but also showed out-of-sample predictive value in held-out participants in a cross-validated model. Multi-region, multi-contrast models based on composite asymmetry measures of PP, PGPe, and SN achieved substantial out-of-sample predictive accuracy. Although DaTSCAN showed higher overall predictive performance, MRI explained additional non-redundant variance beyond DaTSCAN. This result indicates that routine MRI contains clinically meaningful information not fully captured by dopaminergic imaging alone and that integrating these modalities can enhance predictive accuracy.

Prediction was driven primarily by a dominant shared component of asymmetry across the three basal ganglia subregions. The first principal component of the multi-regional asymmetries indicated that the most generalizable clinical signal is largely captured by a common asymmetry pattern shared across these regions. This finding is consistent with a shared pathological process affecting all three regions, whereas region-specific deviations are small and more heterogeneous across subjects. This further suggests that motor symptom lateralization in early PD reflects a shared, circuit-level process across interconnected sensorimotor regions in the basal ganglia^2,22^.

An important limitation of the prediction analysis is that the MRI predictors were pre-selected based on outcome associations within the same cohort. Accordingly, these analyses establish conditional predictive validity of the identified ROI asymmetry signals but do not provide an unbiased estimate of feature selection or predictive performance. Further validation in an independent dataset is key for determining the generalizability of the multi-region MRI asymmetry model in new cohorts.

In conclusion, spatially informed analysis of harmonized structural MRI reveals subregional basal ganglia gradients and hemispheric asymmetry related to PD and motor laterality. By detecting and predicting motor lateralization using gradient-defined posterior “hotspots” and multi-contrast asymmetries, our findings demonstrate that routine MRI can capture spatially dependent signatures of basal ganglia pathology. These results highlight early vulnerability of the sensorimotor nigro-striatopallidal circuit and support a network-level account of motor dysfunction in PD. This framework advances the development of spatially resolved MRI biomarkers for patient stratification and for studying disease mechanisms and longitudinal progression in PD.

## Methods

### Participants

Data used in the preparation of this article were obtained from the Parkinson’s Progression Markers Initiative (PPMI) database (www.ppmi-info.org/access-data-specimens/download-data), RRID:SCR_006431. For up-to-date information on the study, visit www.ppmi-info.org. PPMI – a public-private partnership – is funded by the Michael J. Fox Foundation for Parkinson’s Research, and funding partners, listed in https://www.ppmi-info.org/about-ppmi/who-we-are/study-sponsors.

From the PPMI database, we obtained 633 records from PD and HC participants who underwent similar MRI scan protocols collected with 3T Siemens Tim Trio systems across several imaging centers. Records were excluded for missing MDS-UPDRS-III data (66 PD records), diagnosis duration at baseline greater than 3 years or symptom duration at baseline greater than 10 years (7 PD records), failure of manual image quality assessment (9 PD and 1 HC records), or absence of at least one MRI contrast (T1w, T2w, or PDw; 24 records). After applying these criteria, 526 imaging records remained for analysis across all available visits. The baseline visit included 196 participants (136 PD, 60 HC).

### MRI Acquisition

MRI data were collected using 3T SIEMENS Trio scanners. T1w images were acquired using MPRAGE generalized autocalibrating partially parallel acquisitions (MPRAGE-GRAPPA) sequence, with 1-mm sagittal slice thickness and 1-mm^2^ in-plane resolution. T2w and PDw images were acquired using a turbo spin echo (TSE) sequence with 3-mm axial slice thickness and 0.94-mm^2^ in-plane resolution, either with or without fat suppression (FS).

### Image Processing and Data Harmonization

All MRI analyses were done in each subject’s T1w space. T2w and PDw images were rigidly registered to the T1w image using 9-degree-of-freedom affine transformations. Registration was performed using FSL’s FLIRT^29^ on brain-masked images generated using SynthStrip^30^. Within T1w space, intensity harmonization was applied on each image independently, using a custom subject-level bias-correction tool (https://github.com/MezerLab/mri_unbias) which estimates the spatial bias field as a second-order 3D polynomial fit within white-matter mask, generated using FreeSurfer (v7.4.1)^31^. Because the polynomial directly approximates the white-matter intensity field, dividing the image by this bias map both corrects the spatial bias and implicitly normalizes the white-matter peak to approximately 1. As a result, brain tissue intensities are brought to a common scale across subjects and visits, improving the comparability of T1w, T2w, and PDw images.

### DaTSCAN (123I-FP-CIT SPECT) Striatal Binding Ratios

Dopamine transporter striatal binding ratio (DAT-SBR) values were obtained directly from the PPMI database. DAT-SBR values represent striatal dopamine transporter binding normalized to an occipital cortex reference region, providing a relative index of specific dopaminergic uptake. PPMI provides regional SBR estimates for the left and right caudate, putamen, and anterior putamen. For comparisons with MRI-derived measures, we used the anterior putamen SBR, as it showed the strongest association with motor symptom asymmetry.

### Motor Asymmetry

Motor dysfunction was assessed through MDS-UPDRS III^32^. At the baseline visit, assessment was done off any parkinsonian medication, while in later visits various medication states were applicable. Motor asymmetry was defined as the sum of the scores for the left body side items minus the sum of the scores for the right body side items:

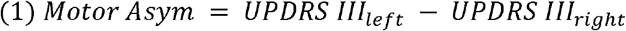

Thus, a greater positive motor asymmetry reflects a left-body laterality and a greater negative motor asymmetry reflects a right-body laterality.

### ROI segmentation

Basal ganglia regions of interest (ROIs) were delineated using a combination of automated segmentation approaches. Caudate nucleus and nucleus accumbens were segmented using FreeSurfer^31^. Putamen was segmented using FSL FIRST probabilistic algorithm^33^, since we observed it had superior anatomical specificity for that region. GP, GPe, GPi, SN, SNpc, SNpr, and STN were segmented using DBSegment^12^. Another, larger, definition of SN was obtained by nonlinearly transforming MASSP subcortical atlas^34^ into each subject’s space, using ANTS^35^. Since the DBSegment SN was smaller than the MASSP SN, and closer to the SN size reported in postmortem literature^36–38^, we termed them SN-small and SN-area, respectively.

For an exploratory analysis, Freesurfer was used to segment the thalamus, amygdala, hippocampus, ventral diencephalon, cerebellar cortex, cerebellar white matter, lateral ventricles, and 31 cortical regions defined by the Desikan-Killiany cortical atlas. Additional subcortical and brainstem nuclei - including the habenular nuclei (HN), red nucleus (RN), ventrointermediate nucleus of the thalamus (VIM), and ventral posterior nucleus of the thalamus (VPL) - were obtained using DBSegment. The ventral tegmental area (VTA), pedunculopontine nucleus (PPN), and periaqueductal gray (PAG) were segmented using the MASSP atlas.

### Putamen and GPe Gradients

AP gradients were quantified along the first major axes of the left and right putamen and GPe at the individual subject level, using the mrGrad tool (https://github.com/MezerLab/mrGrad), as described previously^10,11^. In both the putamen and GPe, the major axes correspond to the regional AP axis. In the putamen, we sampled the MRI T1w, T2w, and PDw intensities in ten equally spaced segments along the AP axis, and in the GPe in five equally spaced segments, considering the regions’ total volume and major axis length.

We restricted gradient analyses to the putamen and GPe. We did not extend this approach to the caudate because prior work in an overlapping early-PD cohort found no PD-related alteration of caudate MRI gradients^10^. Gradient analysis was also not applied to smaller basal ganglia nuclei, including the GPi, SN, SNpc, SNpr, nucleus accumbens and STN, because their small volumes relative to the available MRI resolution (0.07-0.52 cm^3^, as shown in **Table 2**) reduce the reliability of estimating a principal anatomical axis that is robust across individuals. In addition, subdividing these nuclei along such an axis would yield very small subregions. These regions were therefore analyzed using a whole-ROI approach.

### MRI asymmetry measurements

For each ROI and each contrast, asymmetry measures were computed as

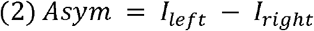

where *I*_*left*_ and *I*_*right*_ are the median signal intensities in the left and right hemisphere ROI, respectively. Volumetric asymmetries were computed as the left mask volume minus the right mask volume in mm^3^.

### Statistical Analysis

#### Gradient spatial and PD-related effects

Statistical inference for spatial and PD-related effects along the major axes of the putamen and GPe was performed using linear mixed-effects models (LMMs), as previously described^10,11^.

For each ROI and MRI contrast, an LMM was fitted with MRI intensity as the dependent variable. Position along the AP axis (interval levels from 0.00 to 1.00) and Research Group (PD, HC categorical levels) were included as fixed effects, together with their interaction. Since age and sex did not differ between groups we did not include them as covariates. Subject ID was modeled as a random intercept to account for within-subject dependence:

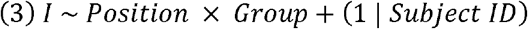

Standardized beta coefficients (*β*) were obtained by z-scoring continuous variables (MRI intensity). Position was not z-scored as it was already normalized to the [0,1] interval, allowing *β*_Position_ to represent the estimated spatial change in MRI intensity along the entire AP axis. Because *Position* was modeled as a continuous predictor, the number of sampling AP nodes did not affect model complexity, but only precision of estimation; this formulation assumes a linear trend between position and the MRI intensity.

The primary effects of interest were the main effect of Position, representing the spatial AP gradient, and the Position × Group interaction, representing group-related alterations of this gradient in PD.

For HC participants and PD participants without motor asymmetry, left and right intensities were averaged before analysis. For PD participants with motor asymmetry, the MAS intensity was used.

##### Post hoc localization

For gradient models showing a significant Position × Group interaction, we performed a post hoc localization analysis. The model was re-estimated with Position treated as a categorical factor, and for each AP position the Group × Position interaction term was tested relative to a reference position (defined as the position with the smallest absolute group difference). To ensure that only positions contributing to the original continuous interaction were retained, the post hoc models computed one-tailed tests based on the sign of the original interaction term. Positions were considered to contribute to the effect at p < 0.01 (uncorrected).

To summarize effects across contrasts and define PD-related subregions, we calculated for each AP position the mean absolute post hoc interaction coefficient across all contrasts, yielding a single cross-contrast interaction magnitude per position. Positions with the largest magnitudes were identified as those exceeding 0.5 robust standard deviations above the median, where the robust standard deviation was estimated from the median absolute deviation (MAD, scaled by 1.4826), a standard robust estimator of σ. Neighboring high-magnitude positions were grouped and treated as subregions, yielding the PP and PGPe subregional ROIs.

#### Simple MRI-motor asymmetry association

Linear regression analyses were performed at the baseline visit to examine the relationship between MRI intensity asymmetries and motor asymmetry. Two families of analyses were conducted. First, basal ganglia regions and subregions (hypothesis-driven ROIs) were tested. Second, exploratory analyses were performed across a broader set of cortical and subcortical regions. Pearson correlation coefficients and associated R^2^ values were computed for each ROI and MRI contrast. Statistical significance was determined using p values adjusted for multiple comparisons across all ROI-contrast combinations within each analysis using the Benjamini-Hochberg false discovery rate method (FDR)^39^.

#### Investigating complementary contributions of MRI contrasts

To examine whether multiple MRI measurements provided complementary information within ROI, we performed multivariate analyses. Within ROI (SN, PP, or PGPe), motor asymmetry was modeled using linear multiple regression model with several MRI measurements. The MRI measurements were selected per ROI based on the single-measurement association results (SN: T2w, PDw, volume; PP: T1w, T2w, PDw; PGPe: T1w, PDw). Thus, these analyses are interpreted as conditional on the prior feature selection.

Complementary value was assessed by comparing the full multivariate model with the best single-measurement model for that ROI using a nested F-test. The resulting ΔR^2^ p values were corrected across the ROI models using FDR.

To assess partial unique contributions of MRI measurements within each ROI model, we examined the coefficient p values from the multivariate regression. These p values were corrected within each ROI across the included MRI measurements using FDR.

To assess whether multiple MRI measurements reflected distinct or shared latent components of clinically relevant variance, we performed PCA within each ROI on the same set of MRI measurements included in the multivariate model. The first principal component (PC1; termed SN_Asym_, PP_Asym_, and PGPe_Asym_) was then tested against motor asymmetry using simple linear regression. Comparison of PC1 and multivariate model performance was descriptive, based on the proportion of variance explained (R^2^_PC1_/R^2^_multi_).

#### Investigating complementary contributions of ROIs

To assess whether different ROIs provided complementary information, we fitted a multi-region multiple regression model at baseline using SN_Asym_, PP_Asym_, and PGPe_Asym_. This ROI set was preselected due to their strong associations with motor asymmetry and anatomical considerations (SN-area, PP, PGPe); therefore, this analysis assessed joint contribution within the selected ROI set rather than across all basal ganglia regions.

We examined the unique contributions of the three ROI predictors within the model and compared the full multi-region model with the best single-ROI model using a nested F-test. Coefficient p values were corrected across the three ROI predictors using FDR.

To assess longitudinal consistency within the cohort, we evaluated the same model at 12, 24, and 48 months using the PCA loadings defined at baseline.

#### Prediction cross-validated modelling

To predict out-of-sample motor asymmetry, we evaluated the multi-region model across all available visits using subject-wise Monte Carlo cross-validation^40^. The MRI predictor set was fixed before cross-validation based on the baseline analyses described above: the three ROI-level MRI components SN_Asym_, PP_Asym_, and PGPe_Asym_. Within each of 30 repetitions, approximately 20% of subjects (all visits) were held out for testing. The model was fitted on the remaining subjects using one randomly selected visit per subject, to avoid dependence among repeated measures in the training set.

In each repetition, normalization and any PCA transformations were estimated on the training data only and then applied to the held-out test data. Importantly, the ROI and MRI-measurement sets themselves were not reselected within cross-validation. Thus, the cross-validated prediction analyses do not reflect unbiased feature selection but rather evaluate conditional predictive validity of the predefined MRI predictors.

We evaluated four cross-validated models: (i) an MRI model using SN_Asym_, PP_Asym_, and PGPe_Asym_; (ii) an MRI PCA model using a single PCA across the selected ROI MRI measurements; (iii) a DaTSCAN-only model using anterior putamen DaTSCAN asymmetry; and (iv) a combined DaTSCAN + MRI model including both DaTSCAN asymmetry and the MRI ROI components.

Out-of-sample predictions were generated for all visits of the held-out subjects. Predictive performance was quantified by the coefficient of determination (R^2^) and root mean square error (RMSE), computed in the test data. We report both the mean ± standard deviation across repetitions and a global estimate obtained by averaging predictions across repetitions for each observation.

To assess the statistical significance of predictive performance, we conducted subject-level permutation tests in which outcomes were randomly re-assigned across subjects while preserving each subject’s visit structure. For each of 500 permutations, the full cross-validation procedure was repeated, and the global predictive R^2^ was recomputed. The permutation p value was defined as the proportion of permuted global R^2^ values greater than or equal to the observed global cross-validated R^2^:

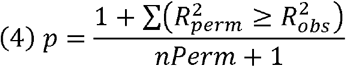

## Author contributions

E.D. and A.A.M. conceived the study, developed the methods, and drafted the manuscript. N.K. performed preliminary analyses. E.D. developed the algorithms, implemented the code, performed the analyses, and prepared the figures.

