## Supplementary Materials for "Sensorimotor basal ganglia circuit asymmetry explains lateralized motor dysfunction in early Parkinson’s disease"

Drori et al. 2026

***Supplementary Table 1. Linear Mixed-Effects Models Result.*** *β coefficients represent standardized response effect estimates. Position effects represent the global change from anterior to posterior. Statistically significant effects after FDR correction are shown in bold.*

| **Putamen T1w** | **β (95% CI)** | **t** | **N** | **DF** | **P** | **P FDR** |
|---|---|---|---|---|---|---|
| **Position** | **3 (2.85, 3.06)** | **55.72** | **196** | **1956** | **0** | **0** |
| ResearchGroup_PD | -0.0068 (-0.16, 0.14) | -0.09 | 196 | 1956 | 0.93 | 0.93 |
| **Position:ResearchGroup_PD** | **-0.13 (-0.26, -0.01)** | **-2.07** | **196** | **1956** | **0.039** | **0.0495** |
| **Putamen T2w** | **β (95% CI)** | **t** | **N** | **DF** | **P** | **P FDR** |
| **Position** | **-2.2 (-2.35, -2.04)** | **-27.62** | **196** | **1956** | **4.1e-142** | **1e-141** |
| **ResearchGroup_PD** | **-0.27 (-0.49, -0.04)** | **-2.35** | **196** | **1956** | **0.019** | **0.028** |
| **Position:ResearchGroup_PD** | **0.45 (0.26, 0.64)** | **4.71** | **196** | **1956** | **2.7e-06** | **6e-06** |
| **Putamen PDw** | **β (95% CI)** | **t** | **N** | **DF** | **P** | **P FDR** |
| **Position** | **-2.5 (-2.66, -2.42)** | **-41.59** | **196** | **1956** | **2.3e-271** | **1e-270** |
| ResearchGroup_PD | -0.15 (-0.36, 0.05) | -1.46 | 196 | 1956 | 0.14 | 0.16 |
| **Position:ResearchGroup_PD** | **0.32 (0.17, 0.46)** | **4.35** | **196** | **1956** | **1.5e-05** | **2e-05** |
| **GPe T1w** | **β (95% CI)** | **t** | **N** | **DF** | **P** | **P FDR** |
| **Position** | **1.5 (1.22, 1.74)** | **11.27** | **196** | **976** | **9e-28** | **2e-27** |
| ResearchGroup_PD | -0.14 (-0.42, 0.15) | -0.95 | 196 | 976 | 0.34 | 0.52 |
| Position:ResearchGroup_PD | -0.13 (-0.44, 0.18) | -0.81 | 196 | 976 | 0.42 | 0.54 |
| **GPe T2w** | **β (95% CI)** | **t** | **N** | **DF** | **P** | **P FDR** |
| **Position** | **2.2 (1.87, 2.43)** | **15.05** | **196** | **976** | **3.3e-46** | **1e-45** |
| ResearchGroup_PD | 0.13 (-0.12, 0.38) | 1.03 | 196 | 976 | 0.3 | 0.52 |
| Position:ResearchGroup_PD | -0.042 (-0.38, 0.29) | -0.25 | 196 | 976 | 0.8 | 0.8 |
| **GPe PDw** | **β (95% CI)** | **t** | **N** | **DF** | **P** | **P FDR** |
| **Position** | **-2.4 (-2.60, -2.18)** | **-22.18** | **196** | **976** | **1.5e-88** | **1e-87** |
| ResearchGroup_PD | 0.033 (-0.21, 0.28) | 0.26 | 196 | 976 | 0.8 | 0.8 |
| **Position:ResearchGroup_PD** | **0.47 (0.21, 0.72)** | **3.60** | **196** | **976** | **0.00034** | **0.00076** |

***Supplementary Table 2. Subregional ROI definition.*** *Subregions are defined on positions where the normalized β coefficient of Position* PD interaction averaged across MRI contrasts was larger than 0.5 (z(β) > 0.5; see* ***Methods****)*

| | **Significant nodes** | **Significant Position** | **z(β) > 0.5 nodes** | **z(β) > 0.5 Position** |
|---|---|---|---|---|
| **Putamen** | 7-9 / 10 | 0.7-0.9 / 1 | 7-9 / 10 | 0.7-0.9 / 1 |
| **GPe** | 3-5 / 5 | 0.6-1 / 1 | 3-4 / 5 | 0.6-0.8 / 1 |

### Supplementary Section 1: Exploratory cortical and subcortical regions asymmetry correlations with motor asymmetry

In addition to the primary basal ganglia ROIs, an exploratory analysis tested associations of MRI intensity asymmetries and motor asymmetry across a broad set of cortical and subcortical regions. Analysis included 46 bilateral regions, including 31 cortical regions, cerebellum cortex, cerebellum white matter, lateral ventricles, thalamus, amygdala, hippocampus, ventral diencephalon, and subregional nuclei: periaqueductal gray (PAG), pedunculopontine nucleus (PPN), ventral tegmental area (VTA), Habenular nuclei (HN), red nucleus (RN), ventrointermediate nucleus of thalamus (VIM), and ventral posterior nucleus of thalamus (VPL).

Several regions showed nominal associations with motor asymmetry (uncorrected $p < 0.05$; **Supplementary Figure 1**); however, none remained significant after correction for multiple comparisons. For context, the Benjamini-Hochberg rank-1 requirement for even a single FDR discovery was $p \leq 0.00036$, corresponding to $R^2 \geq 0.091$. Nominal associations were observed in the VTA (PDw: $R^2 = 0.06$; T2w: $R^2 = 0.03$), PPN (T2w: $R^2 = 0.03$), cerebellum cortex (PDw: $R^2 = 0.03$), lateral occipital cortex (T2w: $R^2 = 0.03$), and pericalcarine cortex (T1w: $R^2 = 0.04$).

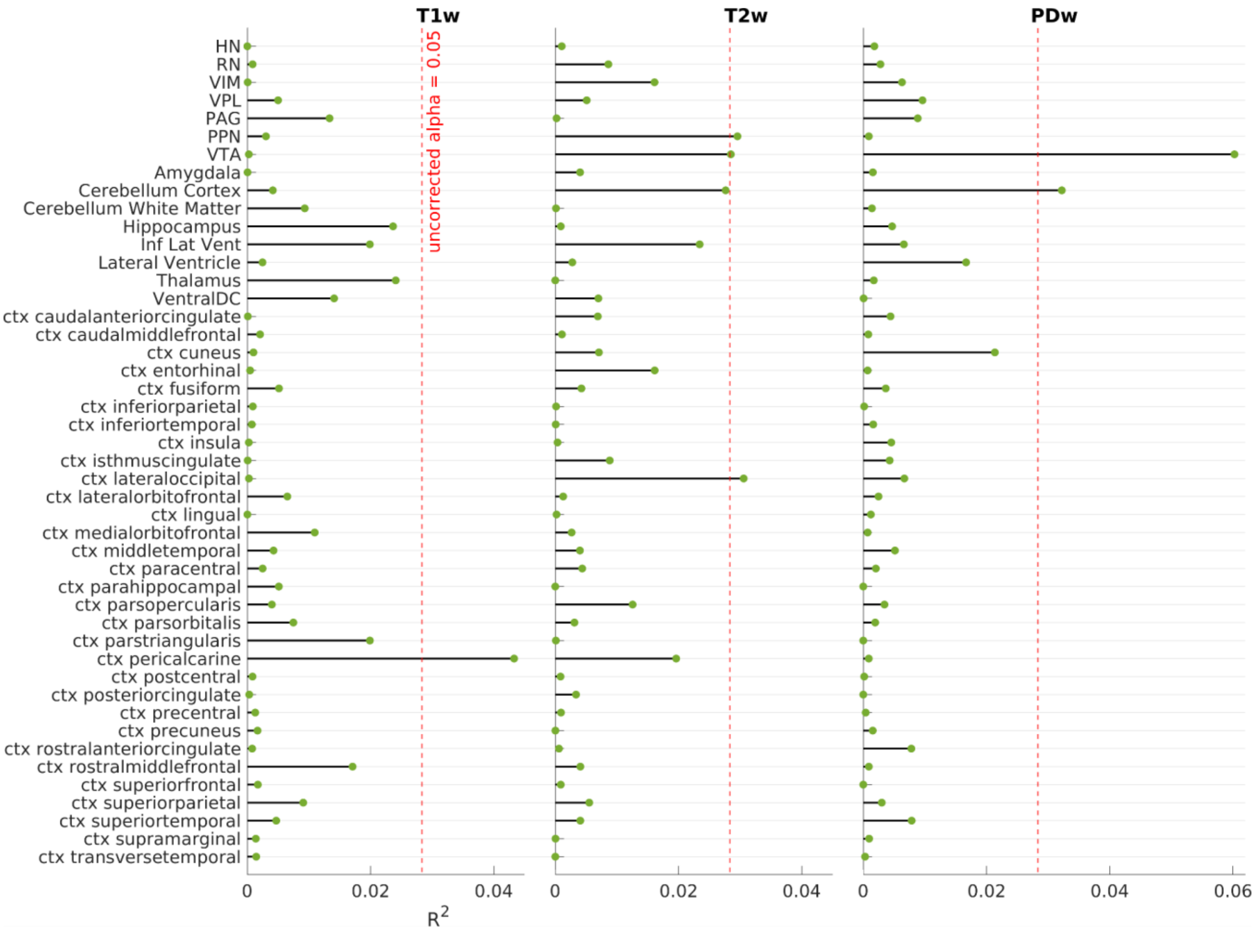


***Supplementary Figure 1. $R^2$ values from associations between exploratory ROIs and motor asymmetry.*** *Forest-style plot displays $R^2$ values of the MRI-motor asymmetry associations across 46 regions. Plot was divided into T1w, T2w, and PDw subplots for convenience. Maximal $R^2$ was 0.06. The red dashed line indicates the uncorrected α = 0.05 critical $R^2$. None remained significant after FDR correction across all comparisons.*

***Supplementary Table 3. MRI contrast contributions in ROI-specific multiple regression models.*** *Multi-contrast models within ROIs (**Figure 4**) show significant partial contributions of each MRI contrast.*

| Model | Measurement | β (95% CI) | SE | t (DF) | partial $R^2$ | P | P FDR |
|---|---|---|---|---|---|---|---|
| **SN** | T2w | 1.96 (0.49, 3.44) | 0.75 | 2.6 (132) | 0.05 | 0.0096 | 0.012 |
| | PDw | 3.00 (1.70, 4.30) | 0.66 | 4.6 (132) | 0.14 | 1.2e-05 | 3.5e-05 |
| | Volume | 1.81 (0.41, 3.22) | 0.71 | 2.6 (132) | 0.05 | 0.012 | 0.012 |
| **PP** | T1w | 1.40 (0.21, 2.60) | 0.60 | 2.3 (132) | 0.04 | 0.022 | 0.022 |
| | T2w | -2.97 (-4.32, -1.62) | 0.68 | -4.3 (132) | 0.13 | 2.7e-05 | 6.5e-05 |
| | PDw | -2.86 (-4.19, -1.52) | 0.68 | -4.2 (132) | 0.12 | 4.3e-05 | 6.5e-05 |
| **PGPe** | T1w | 2.84 (1.54, 4.14) | 0.66 | 4.3 (133) | 0.12 | 3e-05 | 3e-05 |
| | PDw | -3.17 (-4.48, -1.87) | 0.66 | -4.8 (133) | 0.15 | 3.7e-06 | 7.5e-06 |

***Supplementary Table 4. ROI contributions in a multi-ROI multiple regression model.*** *Multi-ROI models (**Figure 5**) show significant partial contributions of multiple ROIs on each visit. FDR-corrected P values above 0.05 are marked in red.*

| Model | Variable | β (95% CI) | SE | t (DF) | Partial $R^2$ | P | P FDR |
|---|---|---|---|---|---|---|---|
| **Baseline** | $SN_{Asym}$ (PC1) | 3.1 (2.07, 4.21) | 0.54 | 5.8 (132) | 0.20 | 4e-08 | 1e-07 |
| | $PP_{Asym}$ (PC1) | -3.3 (-4.63, -2.00) | 0.67 | -5 (132) | 0.16 | 2e-06 | 3e-06 |
| | $PGPe_{Asym}$ (PC1) | -1.8 (-3.11, -0.53) | 0.65 | -2.8 (132) | 0.06 | 0.006 | 0.006 |
| **Month 12** | $SN_{Asym}$ (PC1) | 2.9 (1.57, 4.27) | 0.68 | 4.3 (102) | 0.15 | 4.3e-05 | 0.0001 |
| | $PP_{Asym}$ (PC1) | -1.6 (-3.07, -0.20) | 0.72 | -2.3 (102) | 0.05 | 0.026 | 0.03 |
| | $PGPe_{Asym}$ (PC1) | -2.6 (-3.90, -1.25) | 0.67 | -3.9 (102) | 0.13 | 0.0002 | 0.0003 |
| **Month 24** | $SN_{Asym}$ (PC1) | 2.6 (1.33, 3.93) | 0.65 | 4 (85) | 0.16 | 0.00012 | 0.0004 |
| | $PP_{Asym}$ (PC1) | -2 (-3.40, -0.51) | 0.73 | -2.7 (85) | 0.08 | 0.0085 | 0.009 |
| | $PGPe_{Asym}$ (PC1) | -2.2 (-3.65, -0.73) | 0.73 | -3 (85) | 0.09 | 0.0037 | 0.006 |
| **Month 48** | $SN_{Asym}$ (PC1) | 2.8 (0.88, 4.81) | 0.97 | 2.9 (44) | 0.16 | 0.0054 | 0.008 |
| | $PP_{Asym}$ (PC1) | -4.1 (-6.30, -1.97) | 1.1 | -3.8 (44) | 0.25 | 0.00039 | 0.001 |
| | $PGPe_{Asym}$ (PC1) | -0.21 (-2.37, 1.95) | 1.1 | -0.19 (44) | 0.00 | 0.85 | 0.8 |

##### *Supplementary Table 5. Comparison between MRI and DaTSCAN in motor asymmetry association at Baseline visit.*

| **Modality** | **ROI** | **$R^2$** | **P value** | **Modality's $\Delta R^2$** |
|---|---|---|---|---|
| **DaTSCAN** | Ant. Putamen | 0.58 | $< 10^{-27}$ | 0.11 |
| **MRI** | SN, PP, PGPe | 0.56 | $< 10^{-23}$ | 0.09 |
| **DaTSCAN + MRI** | | 0.67 | $< 10^{-30}$ | Shared Variance: 0.47 |
